# The effects of gamma-aminobutyric acid (GABA) on working memory and attention: A randomised, double-blind, placebo-controlled, crossover trial

**DOI:** 10.1101/2022.11.09.515792

**Authors:** Ahmet Altınok, Aytaç Karabay, Joost de Jong, Gülşen Balta, Elkan G. Akyürek

## Abstract

**Background:** γ-aminobutyric acid (GABA) is a primary inhibitory neurotransmitter that plays a significant role in the central nervous system. Studies on both animals and humans show it has the pharmacological potential for reducing the impact of cognitive disorders, as well as enhancing cognitive functions and mood. However, its specific effects on human attention and working memory have not yet been extensively studied.

**Aims:** In this randomised, double-blind, placebo-controlled, and crossover trial, we aimed to test whether the administration of 800 mg GABA, dissolved in a drink, acutely affected visual working memory maintenance, as well as temporal and spatial attention in healthy adults.

**Methods:** The participants were 32 young adults (16 females and 16 males). Working memory recall precision, spatial attention and temporal attention were measured by a delayed match-to-sample task, a visual search task, and a speeded rapid serial visual presentation task, respectively. Participants completed two experimental sessions (GABA and Placebo) in randomized and counterbalanced order. In each session, forty-five minutes after administration of the drink, they completed the all three of the aforementioned cognitive tasks.

**Results:** Linear mixed model analysis results showed that GABA increased visual search time, compared to the placebo, but did not affect visual search accuracy, temporal attention, nor visual working memory precision.

**Conclusions:** The results suggest that GABA increases visual search time but does not affect temporal attention and memory, and that previously reported effects on cognition might rely on other functions.

## INTRODUCTION

Humans seem to have an innate desire to enhance their abilities, physical as well as cognitive. This is why many people take dietary supplements on a regular basis. One such supplement that has been popular in recent years is gamma-aminobutyric acid (GABA). GABA is an inhibitory neurotransmitter, and can also be found to occur naturally in various foods such as tomatoes, sweet potatoes, spinach and soybeans. Therefore, even without supplementation, many people already consume GABA daily (Briguglio et al., 2018; Ramos-Ruiz et al., 2018). It is conceivable that GABA consumption may have (positive) effects on neuronal signalling, and thereby improve cognition.

Due to its concentration in the central nervous system (CNS), GABA plays a critical role in all mammals (Florey, 1991; Roberts, 2000). In the CNS, GABA acts via its two primary receptors, GABA_A_ and GABA_B_, which are responsible for different regulation systems and include several subtypes. GABA_A_ receptors regulate mood, vigilance, muscle tension and memory functions, while GABA_B_ receptors regulate cerebral reward processes and behaviour (Rudolph & Möhler, 2004; Simeone et al., 2004). Whereas primarily GABA_A_ receptors are associated with cognitive functions, it has been suggested that GABA_B_ receptors also play a crucial role in memory, learning and neurological disorders (Heaney & Kinney, 2016). In particular, subtype α_5_ GABA_A_ receptors are central to the pharmacological modulation of learning and memory. These receptors thus seem crucial to the effect of GABA on cognitive functions (Rudolph & Möhler, 2012; Zheng et al., 2007).

GABA also seems to have region-specific effects within the brain. It has been shown that a decrease in hippocampal GABA_A_ receptor α_5_ subunits disrupts object recognition and memory in mice (Prut et al., 2010). Similarly, reduced hippocampal GABAergic inhibition in GABA_A_ receptors causes lower memory and attention performance in rats (McGarrity et al., 2017). In the prefrontal cortex, GABA activations and levels also influence cognitive functions directly and indirectly. Bañuelos et al. (2014) reported that dysregulation of GABAergic activity in the prefrontal cortex of aged rats negatively affected their working memory performance. Further, Auger and Floresco (2015) showed that antagonism in GABA_A_ signalling in the prefrontal cortex affects search behaviours that are guided by spatial working memory, short-term and reference memory in rats. Nevertheless, the same study showed that blocking GABA’s action does not affect spatial discrimination performance. In other words, GABA’s action did affect working memory performance, but not to the extent that it had a noticeable effect on the precision of memory.

In humans, the prefrontal cortex is prime target for GABAergic interactions, since it is a critical area for executive control (Michels et al., 2012), attention to novel events (Daffner et al., 2000), spatial attention (Small et al., 2003), temporal attention (Coull, 2004), and working memory (Kane & Engle, 2002). Yoon et al. (2016) showed that with a higher working memory load, individuals with lower GABA levels in the prefrontal cortex show a greater reduction in working memory performance from lower loads than participants with higher GABA levels. Notably, the authors suggested that with a high memory load, working memory performance could depend on current GABA levels in the prefrontal cortex and that individual differences in GABA levels could be an important factor for predicting working memory performance. In an fMRI study, Michels et al. (2012) investigated fluctuations in GABA levels in the prefrontal cortex during a Sternberg working memory task in healthy young adults. They observed that GABA levels increase at the outset of the working memory task, and then subsequently decrease as the task progresses. Decreasing GABA levels correlated with lower reaction times and higher levels of accuracy. In other words, GABA’s action on working memory tasks is dynamic.

In addition to the prefrontal cortex, it has been shown that GABAergic actions in the visual cortex have an important role in visual perception (Cook et al., 2016; Yoon et al., 2010). Hammett et al. (2020) tested how differences in GABA concentrations relate to visual contrast discrimination and found that GABA levels positively correlate with visual discrimination performance. Marsman et al. (2017) tested the relationship between the ratio of GABA to glutamate (a dominant excitatory neurotransmitter in the CNS) in the frontal and occipital cortex and the working memory subscale of the Wechsler Adult Intelligence Scale, used extensively to measure general intellectual abilities. The working memory subscale index refers to being able to attend to the information given verbally and the ability to manipulate the information and reproduce it in a short time. Marsman and colleagues found that GABA levels did not correlate with the working memory index, in either the frontal or the occipital cortex, but a higher working memory index did correlate with a lower GABA to glutamate ratio in the frontal cortex, and lower glutamate levels in the occipital cortex. These studies suggest that not only levels of GABA, but also its balance with other neurotransmitters play a role in its functioning.

Even though direct and indirect evidence shows a relationship between GABAergic action and cognitive functions, and its benefits on mood have also been relatively well documented (Brambilla et al., 2003; Petty, 1995; Streeter et al., 2018; Taylor et al., 2003; Yamada et al., 2003; Yoto et al., 2012), there are limited results from randomised control trials testing the effects of GABA ingestion on cognitive functions in humans. This paucity may be related to the idea that GABA cannot pass the blood-brain barrier (Van Gelder & Elliott, 1958, but see also Constans et al., 2020). Nevertheless, some studies have reported effects of GABA consumption on cognition, suggesting that the brain might be affected after all. In young adults, it has been shown that ingestion of 800mg of GABA facilitates temporal attention. Leonte et al. (2018) tested its acute effects on spatial attention using a visual search task and temporal attention, measured by an RSVP attentional blink task. In the visual search task, participants were asked to find a target among non-target items, including different orientations or colours with target items, while the RSVP task was an integration task with Lag1, 3 and 8, without a time limit for responses, as is common in attentional blink research. Results showed that GABA ingestion alleviated temporal attentional bottlenecks at Lag 3; however, it did not affect visual search accuracy and reaction time. However, visual search accuracy ranged from 93.8% to 96%, depending on the task conditions. This might have limited the sensitivity of this task.

Further positive effects of GABA ingestion were reported by Okita et al. (2009), who showed that vegetable tablets containing 30mg GABA acutely benefit the autonomic nervous system in healthy young adults. Okita et al. measured several physiological indicators such as heart rate (HR), heart rate variability (HRV), and ratio of low frequency (LF) and high frequency (HF) of sympathetic and parasympathetic nerve activity, before and after GABA intake. It was reported that tablets acutely decrease the HR and LF/HF ratio. However, in this study, compared to the placebo, the GABA tablets had, in addition to 30mg of GABA, a considerable amount of potassium (520mg), magnesium (50mg), vitamin E (1.49mg), and calcium (232mg).

Finally, Lim and Aquili (2021) tested the acute effects of 800mg of GABA on cognitive flexibility in healthy young adults, using a Stroop task and a task-switching task. They showed that GABA intake decreases cognitive flexibility during task switching, but there was no effect on Stroop task accuracy. The results thus suggested that GABA intake might affect response inhibition, albeit in a negative way. It was suggested that there might be a non-linear, or U-shaped relationship between GABA intake and cognitive performance.

In light of the research to date, it can be said that there are likely relationships between GABA and cognitive functions, and that it has potential for pharmacological interventions for both mood and cognitive disorders. However, the effects of its acute administration on human cognition remain overall not well documented. Although some acute effects on cognition have been observed previously, the studies to date have either tested just one function (e.g., attention), or used tasks that conflated multiple cognitive functions simultaneously. Hence, it remains unclear precisely which functions are affected by GABA and to what degree. In our study, we examined the acute effects of GABA supplementation, on the well-defined cognitive functions of selective attention (temporal and spatial) and working memory. The study was conducted using a crossover, randomised, placebo-controlled, gender-balanced, double-blinded, and counterbalanced design.

## METHOD

### Participants

This study was run with 32 (16 females, 16 males) young adults, aged 18-25 (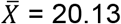, S = 1.81), who were all undergraduate students. A power analysis was run using G*Power software (Faul et al., 2007) for the *F* test, effect size (*f)* = 0.52; α = 0.05; sample size = 16; critical *F* = 4.60. The effect size was calculated based on previous studies (Leonte et al., 2018). Leonte et al. (2018) reported acute effects of GABA on temporal attention with high effect size, 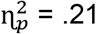 = .21. Thus, a sample size of 32 (16*2 groups/sessions) was needed in total.The following exclusion criteria were set: (1) Not previously diagnosed with any vascular disease, including heart disease (or a history of stroke), hypertension, and diabetes; (2) not previously diagnosed with any health disorder affecting metabolism, such as kidney, liver or gastrointestinal disease; (3) not previously diagnosed with any neurological or psychiatric disorder, including anxiety, depression, migraine and epilepsy; (4) not following a medically restricted diet or taking either over-the-counter or prescription medication, with the exception of the contraceptive pill; (5) not taking vitamin supplements, herbal extracts or illicit drugs; (6) not pregnant or breastfeeding; (7) normal or corrected-to-normal visual acuity, and normal ability to perceive colour and (8) a BMI between 18.5 and 24.9 – defined as the healthy weight range according to the National Health Service of the UK (2019).

### General apparatus

Data was collected in the behavioural laboratories of the Psychology Department of the University of Groningen. A 22” CRT monitor with a refresh rate of 100 Hz and 32-bit colour depth was used. Screen resolution was set to 1280 by 1024 pixels. The experimental tasks were programmed with OpenSesame 3.3.9 (Mathôt et al., 2012), and run under the Windows 10 operating system. Responses were collected via a USB keyboard and mouse.

### General procedure

Participants were recruited via the university participant platform (SONA) and Psychology (paid) Participant Platform, and they either got paid 34 euros or received 5 SONA course credits. In the advertisement for the study, all exclusion criteria (listed above) were shown in detail. Additionally, the experimental process was explained briefly (tasks, drinks, payment/credits, lab visit times etc.). After their enrolment via the participant platforms, they received an email explaining the experimental procedure in detail. Participants were invited to come 45 minutes before the start of the actual experiment to consume one of the experimental drinks. Prior to doing so, they read and signed the informed consent form.

Across two sessions, participants consumed two experimental ingredients dissolved in 200 ml orange juice. Under the GABA condition, this consisted of 800 mg of GABA supplement (from NOW Foods, ordered online at https://www.nowfoods.com/supplements/gaba-powder), and in the placebo condition, this was 800 mg of microcrystalline cellulose. The dosage matched that of a previous study from our group exactly (Leonte et al., 2018). Participants were served these drinks in randomized but counterbalanced order. For instance, if the GABA drink was scheduled on their first visit, they would be served a placebo drink on their second visit or vice versa. The experimental structure is shown in Figure 1.

**Figure 1.**
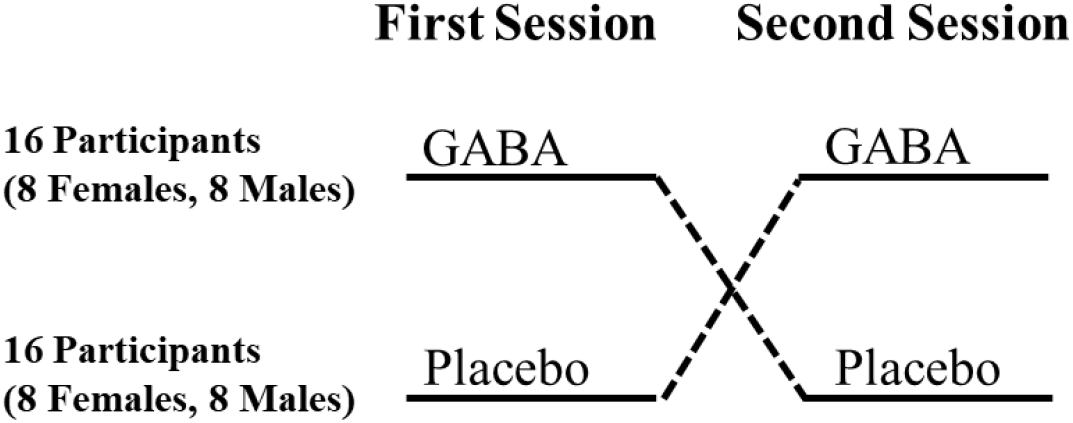
Experimental procedure. Session and drink order was randomized and counterbalanced between equal groups of male and female participants.

After forty-five minutes, when the drinks should have been digested, participants participated in three computerized experimental tasks, which measured temporal attention, spatial attention, and visual working memory. The task order was randomized and counterbalanced for each gender separately. The participants were seated in individual, sound-attenuated testing cabins at approximately 60 cm viewing distance from the screen. During their lab visit, a second researcher was managing the session, to ensure the double-blind procedure. Participants had time to read the instructions of the tasks and to ask questions. They filled out a short questionnaire on the computer just before the experimental tasks, to specify their gender, age, daily caffeine intake, body weight and height. There was at least one week washout period between each session (Leonte et al., 2018), so that there should be no carry-over effects from one session to the next.

### Tasks

As shown in Figure 2, a speeded rapid serial visual presentation (RSVP) task was used to assess temporal attention. A visual search (VS) task was used to measure spatial attention. A delayed match-to-sample task was used to test visual working memory (VWM).

**Figure 2.**
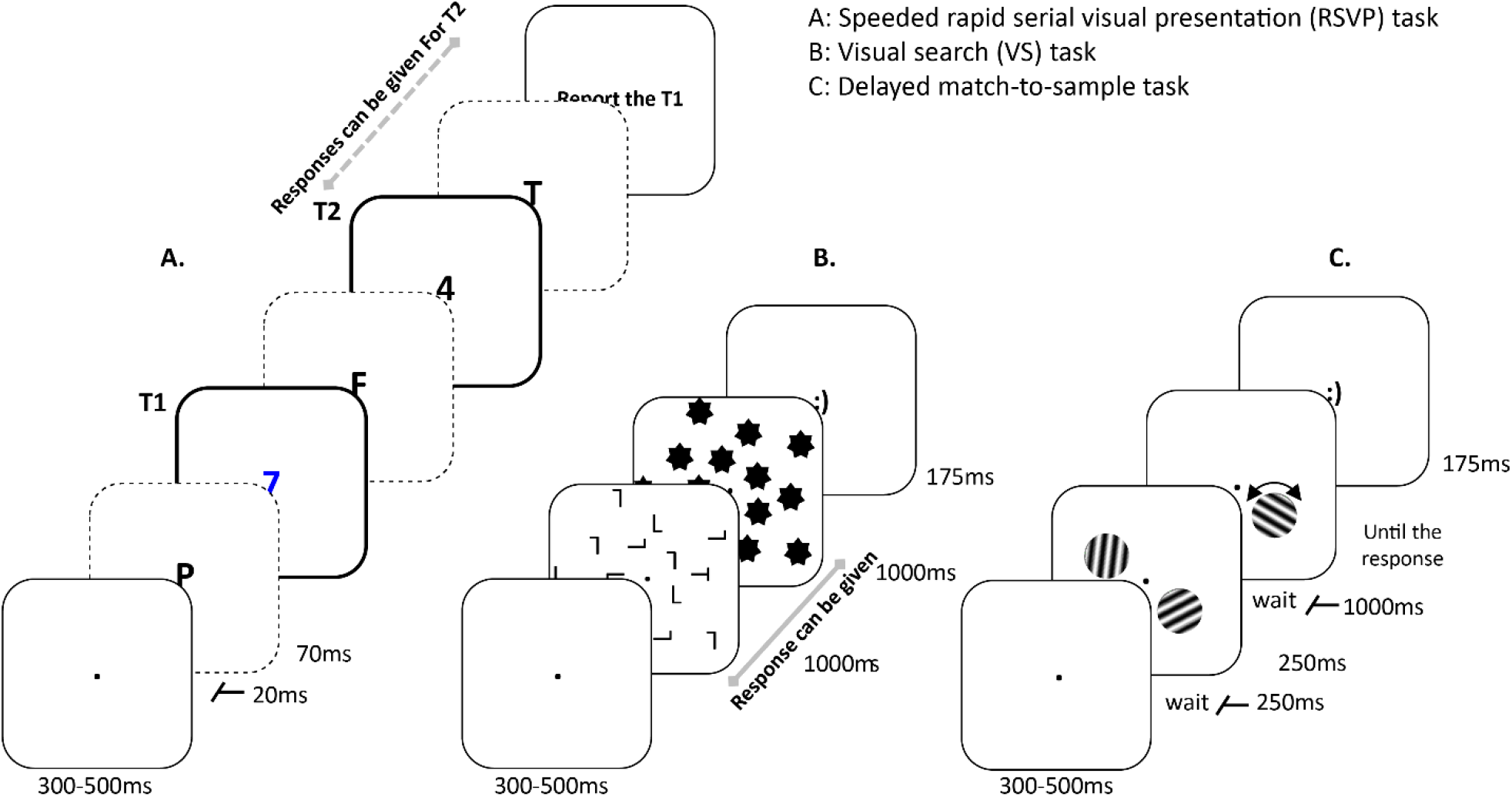
Trial procedures of the experimental tasks. A. An illustration of the speeded RSVP task. Frames with thick outlines indicate the target items (numbers), and dashed outlines indicate a variable numbers of distractors (letters). B. The VS task with 14 distractors. C. The visual working memory task for the two-item condition.

#### Rapid Serial Visual Presentation task

##### Stimuli

A total of 18 stimuli was shown on a light grey background (RGB 192, 192, 192) in RSVP. Targets were numbers between 1 to 9, and distractors were 20 uppercase letters excluding I, O, Q, S, W, and X. The targets and distractors were presented in 52 pt. mono font in the centre of the screen. Target 1 (T1) was presented in blue (RGB 0, 0, 255), Target 2 (T2) and the distractors were presented in black (RGB 0, 0, 0).

##### Procedure

In addition to 30 practice trials, there were 300 experimental trials in the task, spread across a total of ten blocks. Participants were allowed to have a break between each block. At the start of each trial, immediately after the spacebar key was pressed, a black (RGB 0, 0, 0) fixation dot, which had a radius of 8 pixels (6 pt. size), was shown in the centre of the screen for 300-500ms, followed by an RSVP stream comprising two targets and 16 distractors. Each item in the RSVP was presented for 70ms, and trailed by a blank 20ms inter-stimulus interval. The temporal position of T1 was varied randomly from the 5^th^ to the 7^th^ item in the stream, which was evenly distributed across conditions. T2 followed T1 either as the second (Lag 2), third (Lag 3) or eighth (Lag 8) item. Participants were asked to report the identity of targets via the numeric keypad of a standard keyboard. The response to T2 was speeded so that participants were asked to report the T2 identity as soon as they saw it, within 1.5 seconds after T2 onset. The identity of T1 was reported at the end of the trial without time pressure. Overall feedback on task performance was provided between blocks.

##### Design and Analysis

The dependent variables (DVs) were T1 accuracy, conditional T2 accuracy (T2|T1), and T2 reaction time (RT) for correct T2 responses in conditional T2 trials (T2|T1). To clarify, conditional T2 performance refers to trials in which the T1 response was correct. Responses faster than 100 ms were discarded from the T2|T1 RT analysis (three trials, which was equal to 0.03 % of trials).

#### Visual Search task

##### Stimuli

A fixation dot, search array, mask and feedback screen were presented successively on a light grey background (RGB 192, 192, 192). The fixation dot was shown in black (RGB 0, 0, 0), with a radius of 8 pixels (6 pt. size). On the search array, the letters, including the target, were distributed evenly across three invisible, concentric circles. The circles had a radius of 100 pixels, 150 pixels and 200 pixels (2.53°, 3.79° and 5.05° degrees of visual angle, respectively), and were centred on the screen. Each of these invisible circles contained either 5, 7, or 9 letters in randomized but equidistant locations. The target was the letter T, and the distractors were the letter L, both of which appeared in black (RGB 0, 0, 0), in 10 pt. size; 28 × 60 pixels). The orientation of the letters was either 0°, 90°, 180° or 270°, which was evenly distributed and drawn randomly on each trial. The masking display contained black (RGB 0, 0, 0) stars (10 pt. size; 28 × 60 pixels) that appeared on each location that contained a letter in the preceding search display.

##### Procedure

The VS task had 30 practice and 300 experimental trials (100 trials for each condition) with ten blocks. Each block had 30 trials, and participants were allowed to have a break between the blocks. The first trial of each block started when the participant pressed the spacebar. The fixation dot was then shown for 300-500ms, followed by the search array, which was presented for 1000ms and covered by a mask for the next 1000ms. The search array always contained a single target letter, and a variable number of distractors, either 14, 20, or 26. Participants were instructed to report the orientation of the target letter (T) as quickly and accurately as possible. The participants had until the offset of the mask, that is, a total of 2000ms, to give their response. Responses were given with the arrow keys on the keyboard. After the response, participants received feedback for 175 ms: Either a happy or an unhappy smiley depending on their accuracy. The next trial started following an intertrial interval of 250-300ms after the offset of the feedback display.

##### Design and Analysis

Similar to the RSVP analysis, a 2 (two consumption types) by 3 (number of distractors: 14, 20, and 26) factorial design was employed. In this task, DVs were accuracy, and reaction time for correct responses. As before, trials with response times lower than 100ms were excluded from the RT analysis (21 trials, which was equal to 0.13 % of trials).

#### Visual working memory task

##### Stimuli

The stimuli appeared on a light grey background (RGB 192, 192, 192). Each of the memory items was shown at 2.75° of visual angle from the black fixation dot in the centre of the screen, which had a radius of 8 pixels (6 pt. size). The memory items consisted of Gabor patches (sine-wave gratings) of 2.2º of visual angle, and a spatial frequency of 1.8 cycles per degree. The memory items were presented on an invisible circle with a diameter of 6.46º of visual angle. The locations of the memory items on the circle were random, but with the following constraints: In the two memory items condition, the items were presented symmetrically across the two sides of the visual field. In the three items condition, the items were located on an equilateral triangle (still on the circle). The feedback screens consisted of a happy or unhappy smiley in white 32 point size and mono font type, in the centre of the screen.

##### Procedure

The task consisted of 30 practice trials, and 300 experimental trials (100 trials for each item condition), spread across 12 blocks (25 experimental trials each). Between blocks, the participants were allowed to have a break, if they wanted. Each trial started with a fixation dot shown at the centre of the screen for 300 to 500ms. Following a 250ms blank interval, the memory array was shown for 250ms, containing either one, two, or three memory items. The orientation of each item (1-180 º) was chosen randomly with equal probability in each trial. After a one-second delay following the memory array, one of the previously shown item locations was probed with the presentation of another grating in a different orientation, which was randomly chosen, but so that it was at least 15° from the actual orientation of the target memory item. Participants were asked to reproduce the corresponding orientation as accurately as possible by adjusting the orientation of this probe grating. Participants did so by moving a cursor controlled by the USB mouse. Response feedback was given after each trial for 175ms. Positive feedback was given if the error was less than 15°, negative feedback was given otherwise. Furthermore, at the end of each block, block-wise feedback of overall task performance was provided.

##### Design and Analysis

Statistical analysis was performed in the same manner as the other tasks, using a 2 (consumption types) by 3 (number of memory items: 1, 2, and 3) factorial design. The DV was accuracy (% correct), which was calculated based on the degrees of error, where accuracy (%) = 100 - [(100*degrees of error)/90]. Trials with premature responses, that is, with less than 100ms probe response time, were excluded from the analysis (nine trials, which was equal to 0.05 % of trials).

### Data availability

The data and analysis scripts are available on the Open Science Framework (https://osf.io/tqyme).

### Statistical Analysis

A multilevel model (MLM) analysis approach was used to test the acute effects of GABA on temporal attention, spatial attention and visual working memory. Model tests were run via nlme and lme4 (Bates et al., 2015) packages in R (R Core Team, 2012). Linear mixed-effects analysis was performed on dependent variables that had scale outputs (working memory maintenance accuracy, response times of T2|T1 and visual search), and generalized linear mixed-effects analysis was performed on binary (1/0) output dependent variables (T1, T2|T1 and visual search accuracies). As fixed effects, the number of items (working memory), the number of distractors (visual search), Lag (attentional blink), Treatment (Gaba/Placebo), and Session (two sessions) were added to the models. Also, random intercepts for each subject and random slopes for the fixed effects were included in the models.

In the model test process, four steps were followed; first the initial model (without fixed effects) was tested with random intercepts for the subject, then fixed effects were added. Following fixed effects analysis, random slopes were added and at the final stage, pairwise differences were tested with the post-hoc Tukey test. In the model comparisons, simpler models were compared to more complex ones based on the Bayesian Information Criterion (BIC), Bayes Factor -BF_10_-(Wagenmakers, 2007), and differences in deviances with a chi-square test. In the exploratory analysis stage, gender and BMI were entered into the final models as fixed effects.

## RESULTS

### Attentional Blink

In the test of the acute effects of GABA on T1 accuracy with generalized linear mixed model (GLMM) analysis, first, random intercepts for subjects were tested. In the null model (without fixed effects), deviance was 14207, and after adding random intercepts, it was 13089. The model significantly improved when random intercepts for each subject [*X*^2^(1) = 1118.1, *p* < .001, *BF*_*10*_ > 100] were included in the model. Model comparisons showed that adding the fixed effects of Lag (Lag 2, 3 and 8) [*X*^2^(2) = 109.46, *p* < 0.001, *BF*_*10*_ > 100] and Session [*X*^2^(1) = 170.49, *p* < 0.001, *BF*_*10*_ > 100] improved the model significantly, but Treatment did not [*X*^2^(1) = .02, *p* = .905, *BF*_*10*_ = .007]. Random slopes for Session [*X*^2^(2) = 2.52, *p* = .774, *BF*_*10*_ > 100] improved the model significantly but Lag did not [*X*^2^(5) = 9.77, *p* = .082, *BF*_*10*_ < .001]. The final model result is shown in Table 1.

**Table 1.** Generalized Linear Mixed Model (GLMM) Results for T1 Accuracy - Final Model

|  | T1 ACC |  |  |  |
| --- | --- | --- | --- | --- |
| Fixed Effects | Odds Ratios | S.E. | z | p |
| (Intercept) | 3.28 | 1.24 | 5.42 | <0.001 |
| Lag [Lag3] | 1.35 | 1.06 | 5.43 | <0.001 |
| Lag [Lag8] | 1.84 | 1.06 | 10.51 | <0.001 |
| Session | 1.91 | 1.12 | 5.82 | <0.001 |
| Random Effects | Variance | Sd |  |  |
| $\tau_{00}$ Subject | 1.27 | 1.13 | | |
| $\tau_{11}$ Subject. Session | 0.28 | 0.53 | | |
| N<sub>subject</sub> = 32, Observations = 20736 |  |  |  |  |

In pairwise comparisons, Tukey test results showed that T1 accuracy at Lag 8 (*prob* = .941, SE = .009) was significantly higher than at Lag 2 (*prob =* .897, SE = .015), Z = 10.51, *p* < .001, and Lag 3 (*prob =* .921, SE = .012), Z = 5.20, *p* < .001. Also, T1 accuracy at Lag 3 was higher than Lag 2, Z = 5.43, *p* < .001 (see Fig. 3). Regarding sessions, as expected, T1 accuracy was higher in the second session (*prob =* .942, SE = .009), compared to the first session (*prob =* .895, SE = .015), Z = 5.82, *p* < .001.

**Figure 3.**
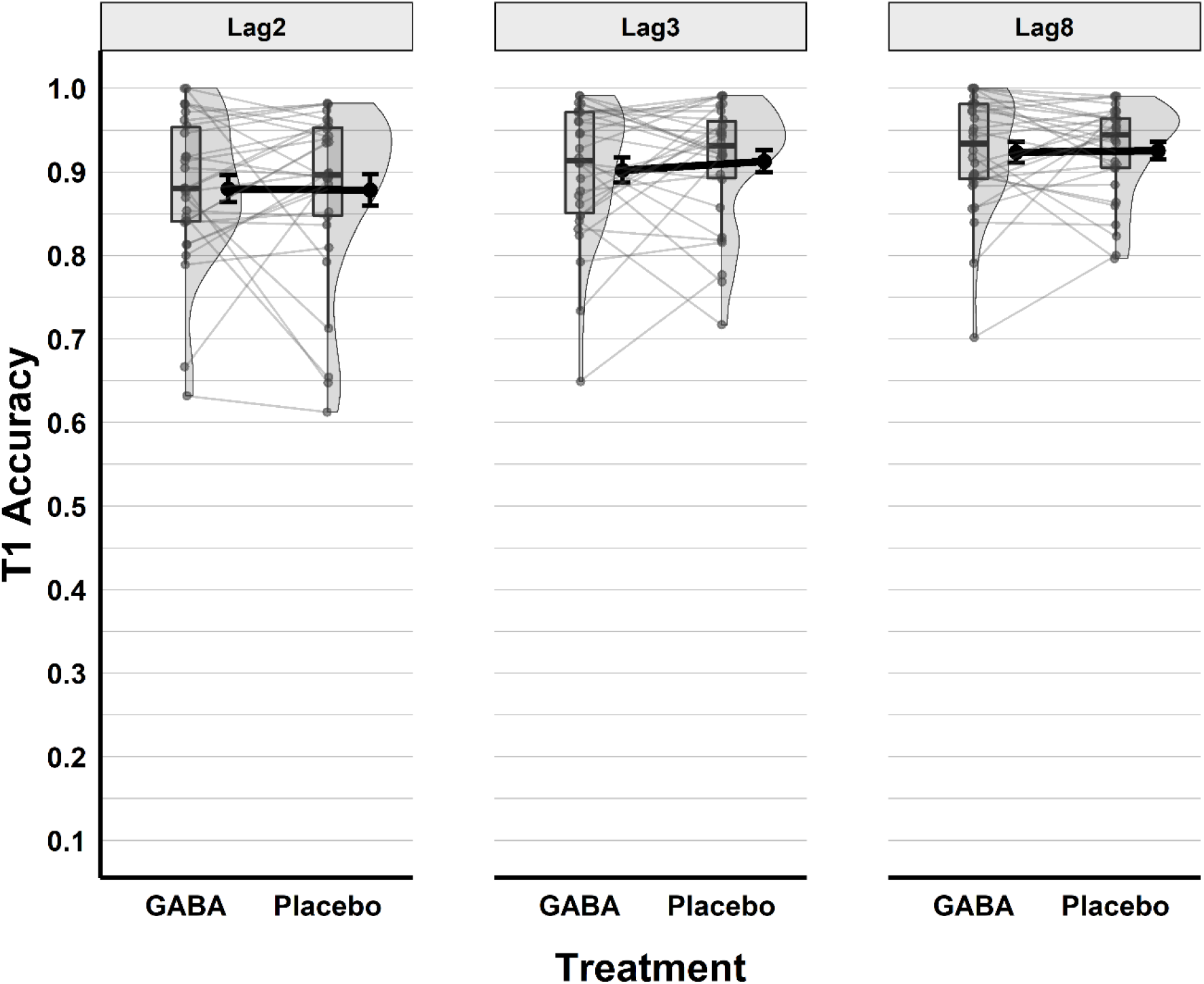
Speeded RSVP T1 accuracy by Treatment (GABA and Placebo) and Lag. Boxplots show quartiles, black dots with error bars show means and standard errors, individual data points are presented with grey points and lines connect individual data points across treatments.

Additionally, in exploratory analyses, the effects of BMI and gender were tested. Analyses results showed that neither gender [*X*^2^(1) = .08, *p* = .772, *BF*_*10*_ = .007] nor BMI [*X*^2^(1) = .74, *p* = .388, *BF*_*10*_ = .010] improved the model at all.

On T2|T1 accuracy, GLMM results showed that random intercepts for each subject [*X*^2^(1) = 1367.8, *p* < .001, *BF*_*10*_ > 100] were necessary. Adding Lag [*X*^2^(2) = 992.92, *p* < .001, *BF*_*10*_ > 100] and Session [*X*^2^(1) = 315.52, *p* < .001, *BF*_*10*_ > 100] as fixed effects improved the model significantly, but Treatment [*X*^2^(1) = 8.31, *p* = .004, *BF*_*10*_ = .468] did not.

Lastly, random slopes for Lag [*X*^2^(5) = 208.9, *p* < .001, *BF*_*10*_ > 100] and Session [*X*^2^(4) = 123.5, *p* < .001, *BF*_*10*_ > 100] were added to the model, and the model improved significantly. The final model results are given in Table 2.

**Table 2.**
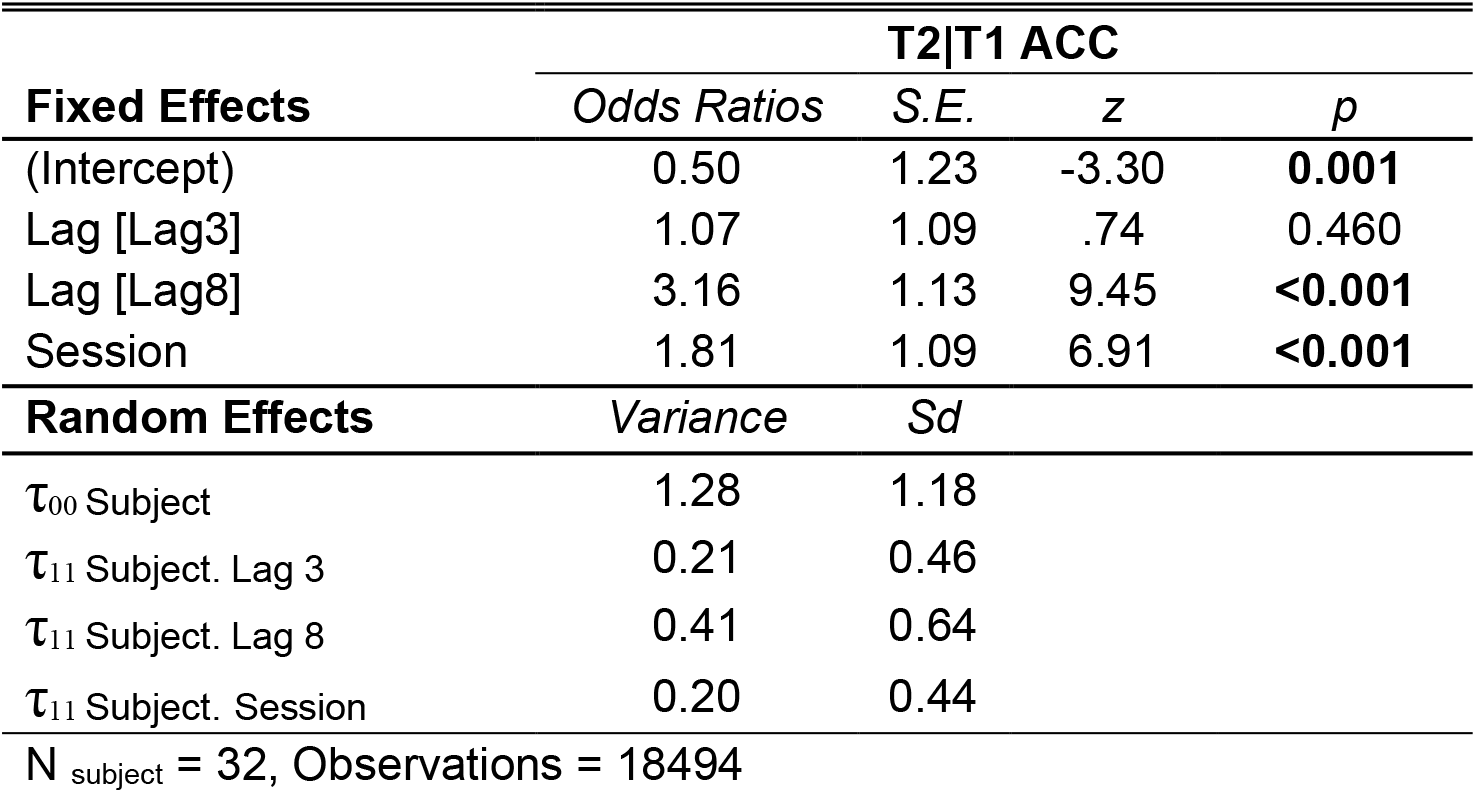
Generalized Linear Mixed Model (GLMM) Results for T2|T1 Accuracy – Final Model

In the post-hoc analysis, Tukey test results showed that T2|T1 accuracy was significantly higher in the Lag 8 condition (*prob =* .794, SE = .021), compared to Lag 2 (*prob =* .549, SE = .036), Z = 9.45, *p* < .001, and Lag 3 (*prob =* .566, SE = .036), Z = 11.99, *p* < .001. However, there were no differences between Lag 3 and Lag 2, Z = .75, *p* = .74 (see Fig. 4). Between sessions, similar to T1, T2|T1 accuracy was higher in the second session (*prob =* .711, SE = .026), than in the first session (*prob =* .577, SE = .035), Z = 6.91, *p* < .001.

**Figure 4.**
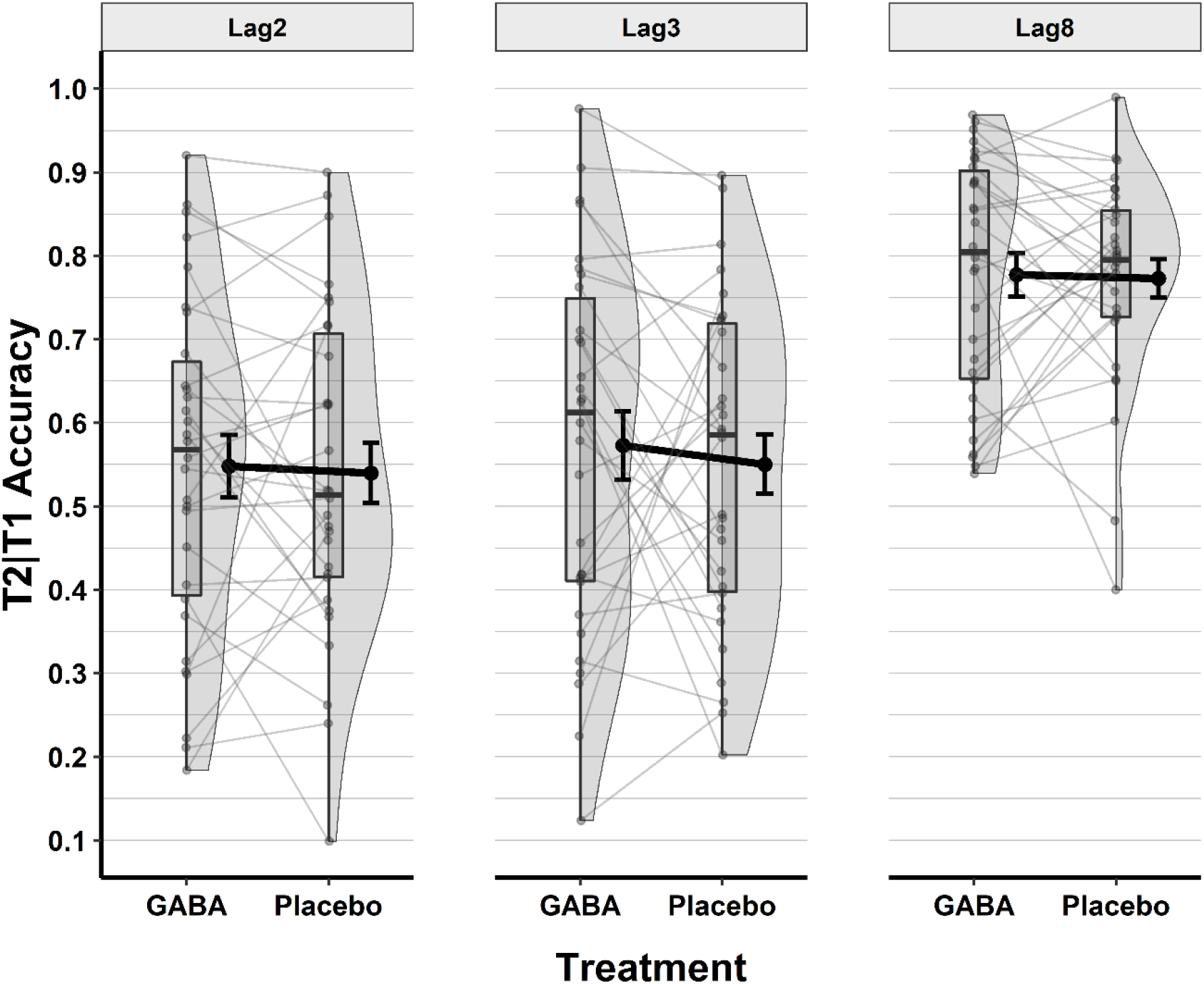
Speeded RSVP T2|T1 accuracy by Treatment (GABA and Placebo) and Lag. Boxplots show quartiles, black dots with error bars show means and standard errors, individual data points are presented with grey points, and lines connect individual data points across treatments.

Finally, neither the gender of participants [*X*^2^(1) = .387, *p* = .534, *BF*_*10*_ = .009], nor their BMI scores [*X*^2^(1) = 3.38, *p* = .066, *BF*_*10*_ = .041] improved the model significantly.

In the RSVP-speeded task, response times for T2 trials were also measured (Fig. 5). The effects of GABA on response times for correct T2|T1 and T1 trials were tested with Linear Mixed Model (LMM) analysis. Adding random intercepts for subjects [*X*^2^(1) = 3232.7, *p* < .001, *BF*_*10*_ > 100] improved the model significantly. Fixed effects analysis results revealed that Lag [*X*^2^(2) = 2388.8, *p* < .001, *BF*_*10*_ > 100] and Session [*X*^2^(1) = 540.13, *p* < .001, *BF*_*10*_ > 100] improved the model significantly but Treatment [*X*^2^(1) = 2.72, *p* = .099, *BF*_*10*_ = .408] did not. Random slopes of Lag [*X*^2^(5) = 308.32, *p* < .001, *BF*_*10*_ > 100] and Session [*X*^2^(4) = 627.16, *p* < .001, *BF*_*10*_ > 100] also improved the model. Table 3 shows the final model results.

**Table 3.** Linear Mixed Model (LMM) Results for T2|T1 Response Time - Final Model

|  | T2\|T1 RT |  |  |  |
| --- | --- | --- | --- | --- |
| Fixed Effects | Estimates | S.E. | t | p |
| (Intercept) | 1268.93 | 56.19 | 22.59 | <0.001 |
| Lag [Lag3] | -99.18 | 10.85 | -9.14 | <0.001 |
| Lag [Lag8] | -283.78 | 18.23 | -15.57 | <0.001 |
| Session | -114.71 | 22.86 | -5.02 | <0.001 |
| Random Effects | Variance | Sd |  |  |
| $\sigma^2$ (Residual) | 51234.32 | 226.35 | | |
| $\tau_{00}$ Subject | 98857.61 | 314.42 | | |
| $\tau_{11}$ Subject. Lag 3 | 2689.80 | 51.86 | | |
| $\tau_{11}$ Subject. Lag 8 | 9715.10 | 98.57 | | |
| $\tau_{11}$ Subject. Session | 16095.19 | 126.87 | | |
| N<sub>subject</sub> = 32, Observations = 11582 |  |  |  |  |

**Figure 5.**
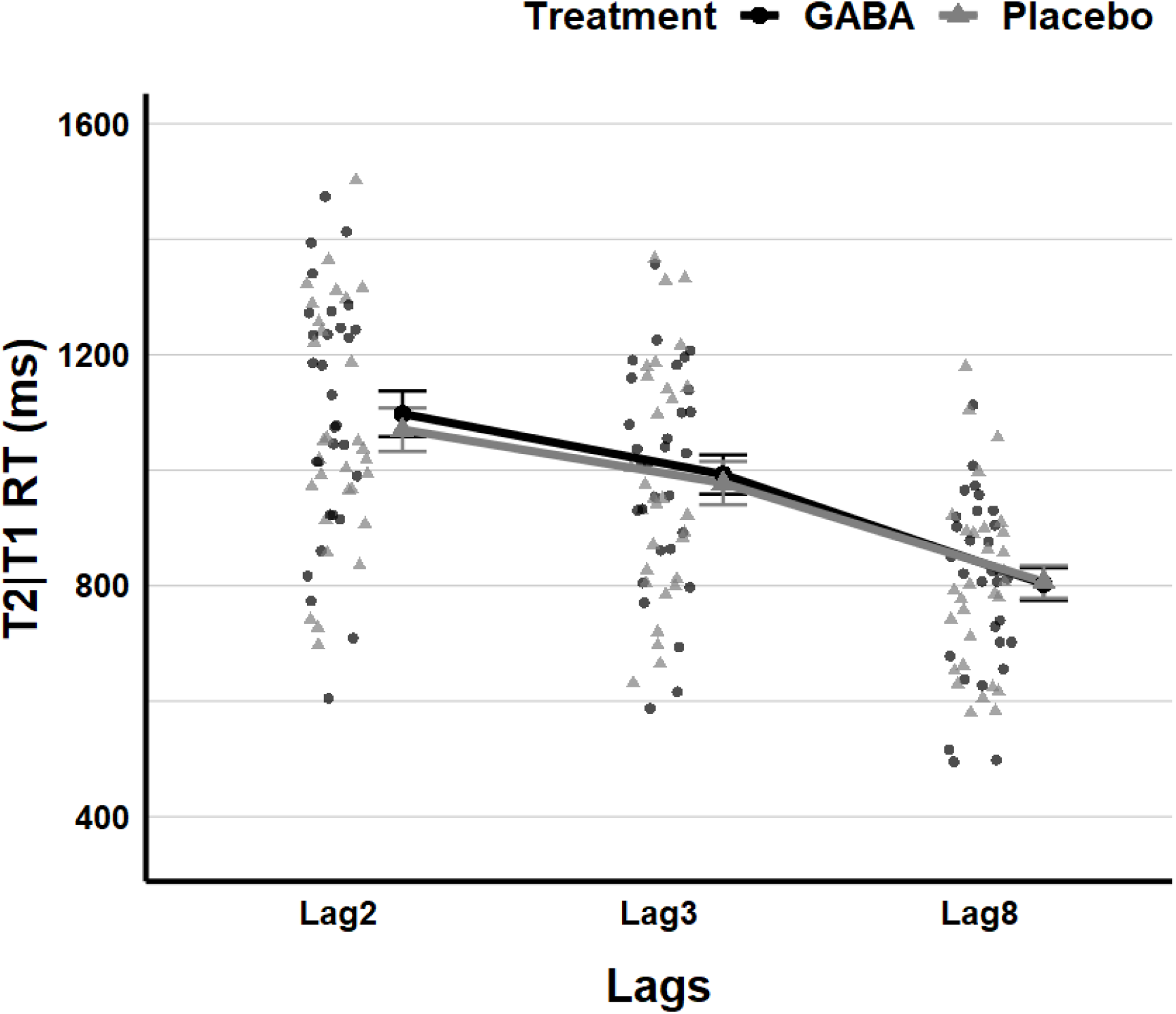
Speeded RSVP T2|T1 response time by Treatment (GABA and Placebo) and Lag. Dots represent individual reaction time means in different treatment conditions. Bigger black dots with error bars show means and standard errors in the GABA condition, while light grey triangles with error bars display means and standard errors in the Placebo condition.

In the post-hoc analysis, Tukey results showed that reaction time at Lag 8 (*Estimated marginal mean =* 813.0, SE = 24.3) was lower than at Lag 3 (*Estimated marginal mean = 998*.0, SE = 32.4), Z = 12.61, *p* < .001, and Lag 2 (*Estimated marginal mean = 1097*.0, SE = 35.7), Z = 15.57, *p* < .001. Also, response time at Lag 3 was lower than at Lag 2, Z = 9.14, *p* < .001. Between sessions, participants were faster to respond to T2|T1 in their second session (*Estimated marginal mean =* 912, SE = 28.2), than they were in the first session (*Estimated marginal mean =* 1027, SE = 35.6), Z = 5.02, *p* < .001.

Lastly, neither the gender of participants [*X*^2^(1) = 1.41, *p* = .234, *BF*_*10*_ = 1.832], nor the BMI scores [*X*^2^(1) = .31, *p* = .578, *BF*_*10*_ = .259] improved the model at all.

### Visual Search

The effects of GABA on visual search accuracy were tested with GLMM analysis. During the model analysis, random intercepts for subjects [*X*^2^(1) = 1003.1, *p* < .001, *BF*_*10*_ > 100] were added to the model. The number of distractors [*X*^2^(2) = 287.22, *p* < .001, *BF*_*10*_ > 100] and Session [*X*^2^(1) = 336.73, *p* < .001, *BF*_*10*_ > 100] were added to the model as fixed effects, which improved the model significantly, but Treatment did not improve the model [*X*^2^(1) = .66, *p* = .416, *BF*_*10*_ = .009]. Adding random slopes for Session [*X*^2^(2) = 26.62, *p* < .001, *BF*_*10*_ = 29.05] significantly improved the model, but random slopes for the number of distractors [*X*^2^(5) = 7.42, *p* < .191, *BF*_*10*_ =.010] did not. The final model results of GLMM analysis for visual search accuracy are shown in Table 4.

**Table 4.** Generalized Linear Mixed Model (GLMM) Results for Visual Search Accuracy - Final Model

|  | VS ACC |  |  |  |
| --- | --- | --- | --- | --- |
| Fixed Effects | Odds Ratios | S.E. | z | p |
| (Intercept) | 2.26 | 1.15 | 5.93 | <0.001 |
| Number of Distractors [20] | 0.62 | 1.05 | -10.65 | <0.001 |
| Number of Distractors [26] | 0.48 | 1.04 | -16.76 | <0.001 |
| Session | 1.91 | 1.06 | 10.87 | <0.001 |
| Random Effects | Variance | Sd |  |  |
| $\tau_{00}$ Subject | 0.48 | 0.69 | | |
| $\tau_{11}$ Subject. Session | 0.07 | 0.26 | | |
| N<sub>subject</sub> = 32, Observations = 20736 |  |  |  |  |

In pairwise comparisons, the Tukey test results showed that search accuracy was significantly higher when 14 distractors were shown (*prob =* .*857*, SE = .014), compared to when either 20 (*prob =* .788, SE = .019), Z = 10.65, *p* < .001, or 26 distractors were presented (*prob =* .742, SE = .021), Z = 16.76, *p* < .001. Additionally, in the 20 distractor condition, search accuracy was significantly higher than in the 26 distractors condition, Z = 6.32, *p* < .001 (Fig. 6). Across sessions, accuracy was higher in the second session (*prob =* .847, SE = .015), compared to the first session (*prob =* .743, SE = .021), Z = 10.87, *p* < .001. As in RSVP, on visual search accuracy, neither the gender of participants [*X*^2^(1) = .59, *p* = .439, *BF*_*10*_ =.009] nor the BMI scores [*X*^2^(1) = 1.501, *p* = .221, *BF*_*10*_ =.015] improved the model.

**Figure 6.**
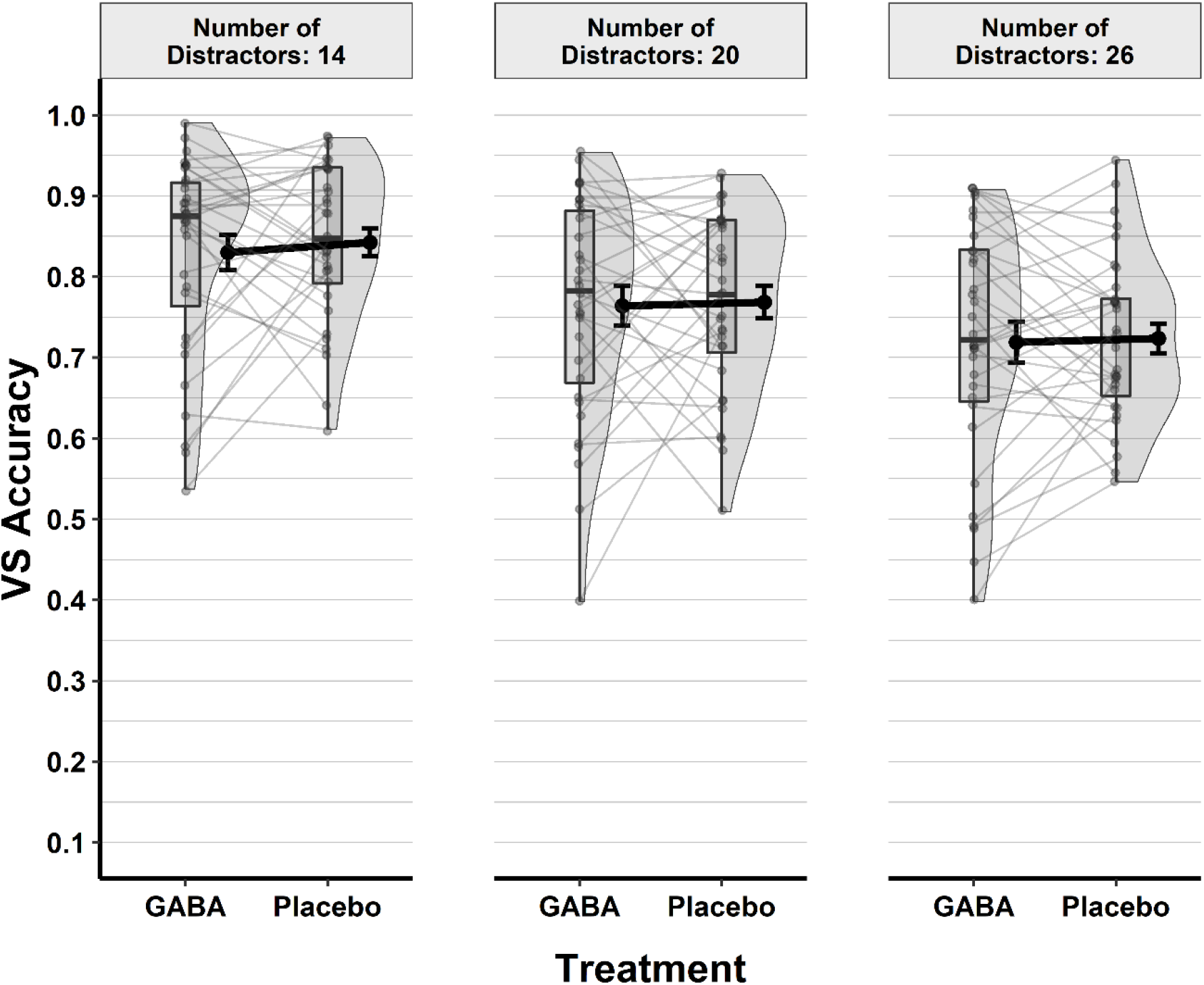
Visual Search accuracy by Treatment (GABA and Placebo) and the number of distractors. Boxplots show quartiles, black dots with error bars show means and standard errors, individual data points are presented with grey points, and lines between individual data points show changes in treatment.

Linear mixed model (LMM) analysis was used to test the effects of GABA on visual search time (Fig. 7). It was revealed that random intercepts for each subject [*X*^2^(1) = 1571.4, *p* < .001, *BF*_*10*_ > 100] were necessary. As fixed effects, the number of distractors [*X*^2^(2) = 302.63, *p* < .001, *BF*_*10*_ > 100], Session [*X*^2^(1) = 619.92, *p* < .001, *BF*_*10*_ > 100] and Treatment [*X*^2^(1) = 57.24, *p* < .001, *BF*_*10*_ > 100] improved the model significantly, while the interaction of Treatment and the number of distractors [*X*^2^(1) = 3.16, *p* = .205, *BF*_*10*_ = .241] did not. Random slopes for Session [*X*^2^(2) = 201.36, *p* < .001, *BF*_*10*_ > 100] also improved the model but neither the number of distractors [*X*^2^(5) = 6.23, *p* = .284, *BF*_*10*_ < .0001] nor Treatment [*X*^2^(3) = 3.19, *p* = .363, *BF*_*10*_ < .0001] did. The final model results are shown in Table 5.

**Table 5.** Linear Mixed Model (LMM) Results for Visual Search Time - Final Model

| Fixed Effects | VS RT |  |  |  |
| --- | --- | --- | --- | --- |
|  | Estimates | S.E. | t | p |
| (Intercept) | 1184.39 | 32.85 | 36.05 | <0.001 |
| Number of Distractors [20] | 67.84 | 5.89 | 11.52 | <0.001 |
| Number of Distractors [26] | 106.69 | 5.99 | 17.83 | <0.001 |
| Session | -121.61 | 15.48 | -7.86 | <0.001 |
| Treatment [Placebo] | -34.43 | 15.44 | -2.23 | 0.026 |
| Random Effects | Variance | Sd |  |  |
| $\sigma^2$ (Residual) | 95639.68 | 309.3 | | |
| $\tau_{00}$ Subject | 30224.64 | 173.9 | | |
| $\tau_{11}$ Subject. Session | 6886.49 | 82.98 | | |

N<sub>subject</sub> = 32, Observations = 16043

**Figure 7.**
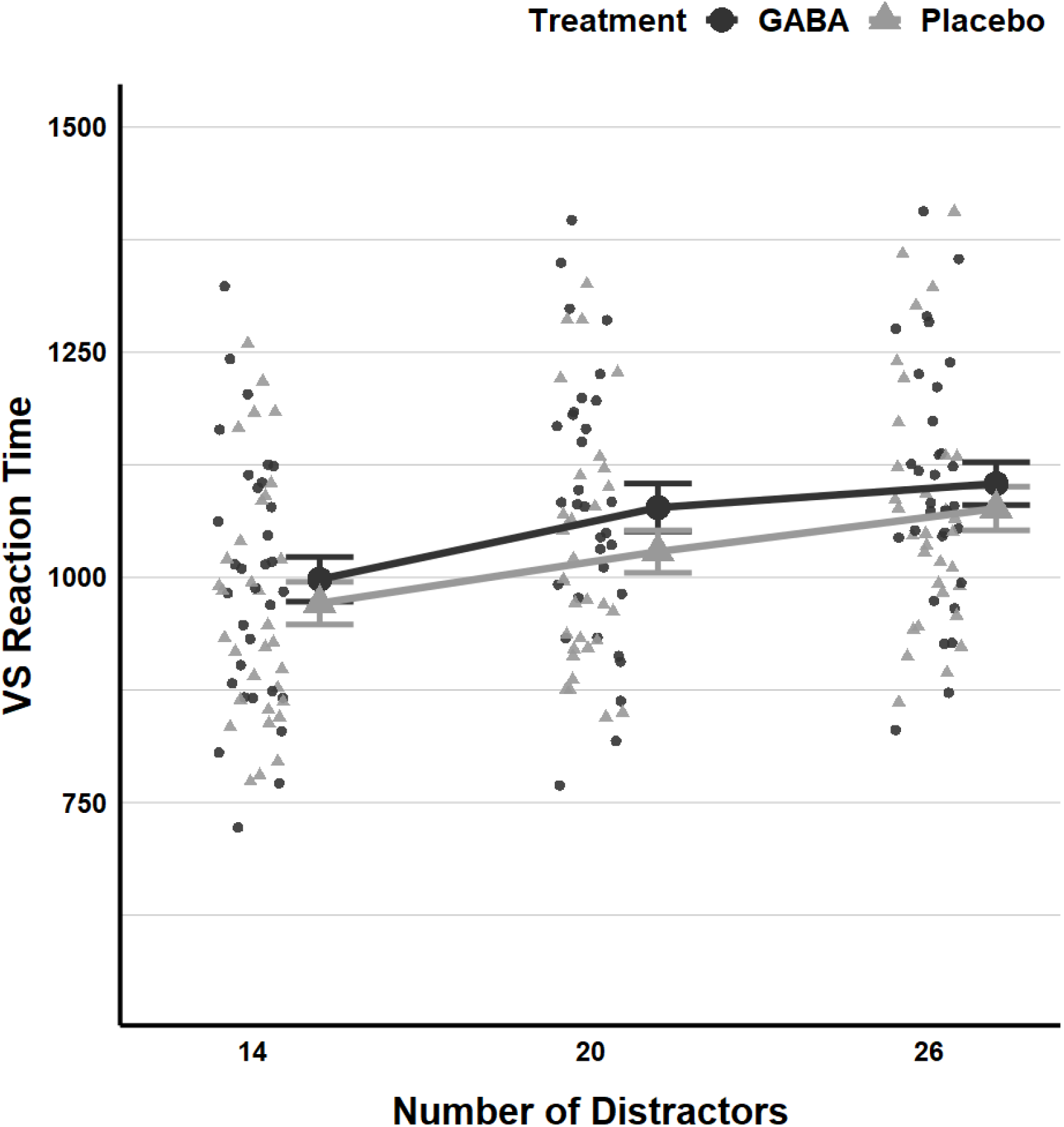
Visual Search reaction time by Treatment (GABA and Placebo) and Lag. Dots represent individual reaction time means in different treatment conditions. Bigger black dots with error bars show means and standard errors in the GABA condition, while light grey triangles with error bars display means and standard errors in the Placebo condition.

Tukey pairwise comparisons showed that visual search time was lower when 14 distractors (*Estimated marginal mean =* 985.0, SE = 20.2) were displayed than when 20 (*Estimated marginal mean =* 1053.0, SE = 20.3), Z = 11.52, *p* < .001, and 26 distractors were displayed (*Estimated marginal mean =* 1091.0, SE = 20.3), Z = 17.83, *p* < .001. Also, in the 20 distractors condition, the search process was faster than in the 26 distractors condition, Z = 6.36, *p* < .001. Similar to search accuracy, visual search time was lower in the second session (*Estimated marginal mean =* 928, SE = 20.9), than in the first session (*Estimated marginal mean =* 1104, SE = 22.0), Z = 7.86, *p* < .001.

Across treatment conditions (See Fig. 8), response times were significantly higher in the GABA condition (*Estimated marginal mean =* 1060, SE = 21.4) than in the placebo condition (*Estimated marginal mean =* 1026, SE = 21.4), Z = 2.23, *p* = .0257. On visual search reaction time, neither the gender of participants [*X*^2^(1) = .27, *p* = .602, *BF*_*10*_ = .904], nor the BMI scores [*X*^2^(1) = .11, *p* = .734, *BF*_*10*_ = .201] improved the model.

**Figure 8.**
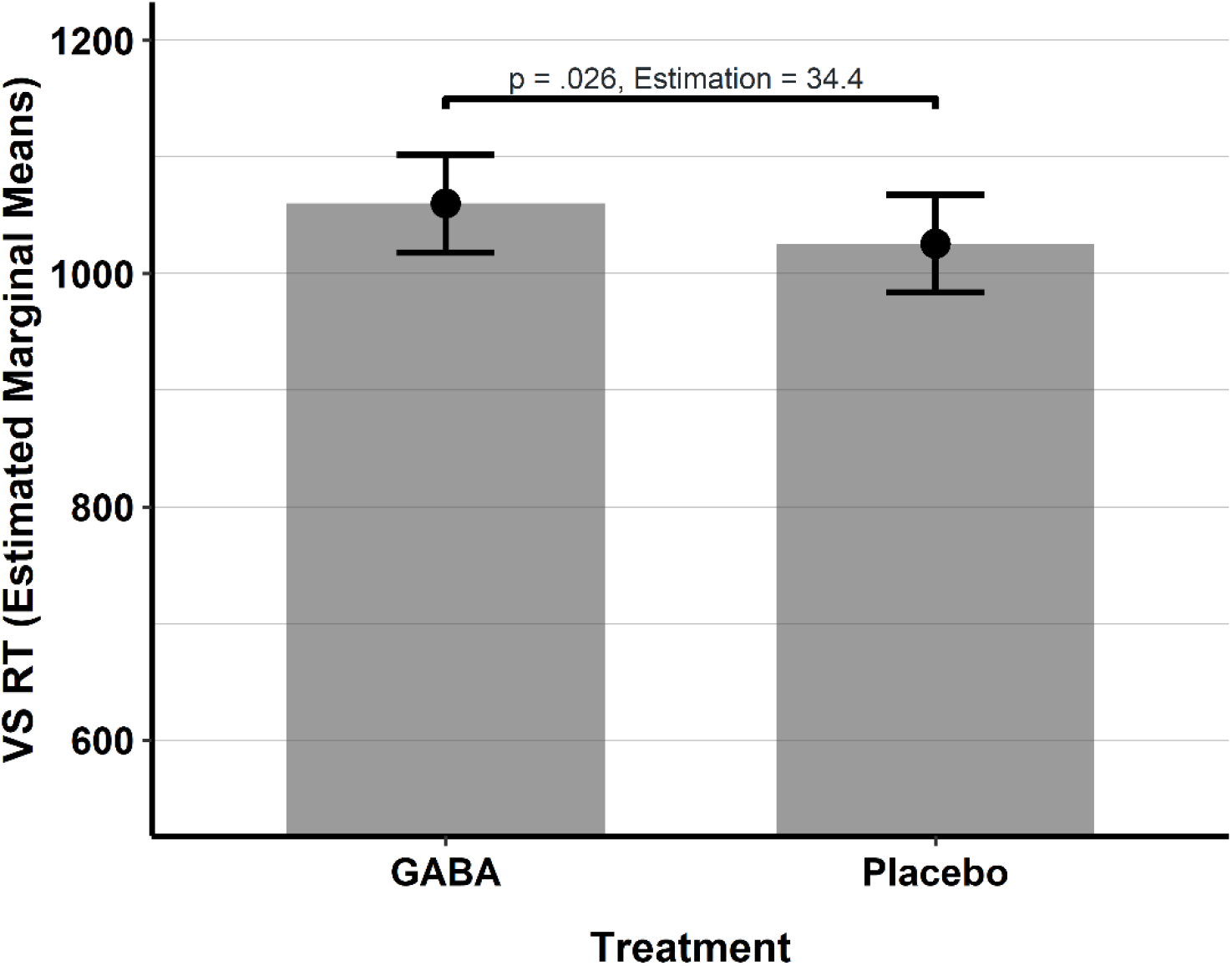
Visual Search reaction time by Treatment (GABA and Placebo). Bar plots with error bars show estimated marginal means and 95 % confidential intervals in Treatment conditions from the final model. *p < .05

### Visual Working Memory

The effects of GABA on working memory were tested with linear mixed model analysis (Fig. 9). Model tests showed that adding random intercepts for each subject [*X*^2^(1) = 1014.5, *p* < .001, *BF*_*10*_ < 100] was necessary. The number of items [*X*^2^(2) = 2509.4, *p* < .001, *BF*_*10*_ < 100] and Session [*X*^2^(1) = 40.2, *p* < .001, *BF*_*10*_ < 100], as fixed effects, improved the model significantly, but the model did not improve when Treatment was included [*X*^2^(1) = 1.971, *p* = .160, *BF*_*10*_ = .014]. Random slopes for the number of items [*X*^2^(5) = 231.46, *p* < .001, *BF*_*10*_ > 100] improved the model, but adding random slopes for Session [*X*^2^(4) = 19.03, *p* = .001, *BF*_*10*_ = .00005] did not. The final model results are given in Table 6.

**Table 6.** Linear Mixed Model (LMM) Results for Working Memory Maintenance - Final Model

|  | WM ACC |  |  |  |
| --- | --- | --- | --- | --- |
| Fixed Effects | Estimates | S.E. | t | p |
| (Intercept) | 88.18 | .62 | 141.12 | <0.001 |
| Number of Items [Three Items] | -17.81 | 1.03 | -17.32 | <0.001 |
| Number of Items [Two Items] | -8.31 | .66 | -12.61 | <0.001 |
| Session | 1.78 | .28 | 6.38 | <0.001 |
| Random Effects | Variance | Sd |  |  |
| $\sigma^2$ (Residual) | 372.79 | 19.31 | | |
| $\tau_{00}$ Subject | 5.04 | 2.44 | | |
| $\tau_{11}$ Subject. Number of Items [Three Items] | 30.10 | 5.49 | | |
| $\tau_{11}$ Subject. Number of Items [Two Items] | 10.16 | 3.19 | | |
| N<sub>subject</sub> = 32, Observations = 19191 |  |  |  |  |

**Figure 9.**
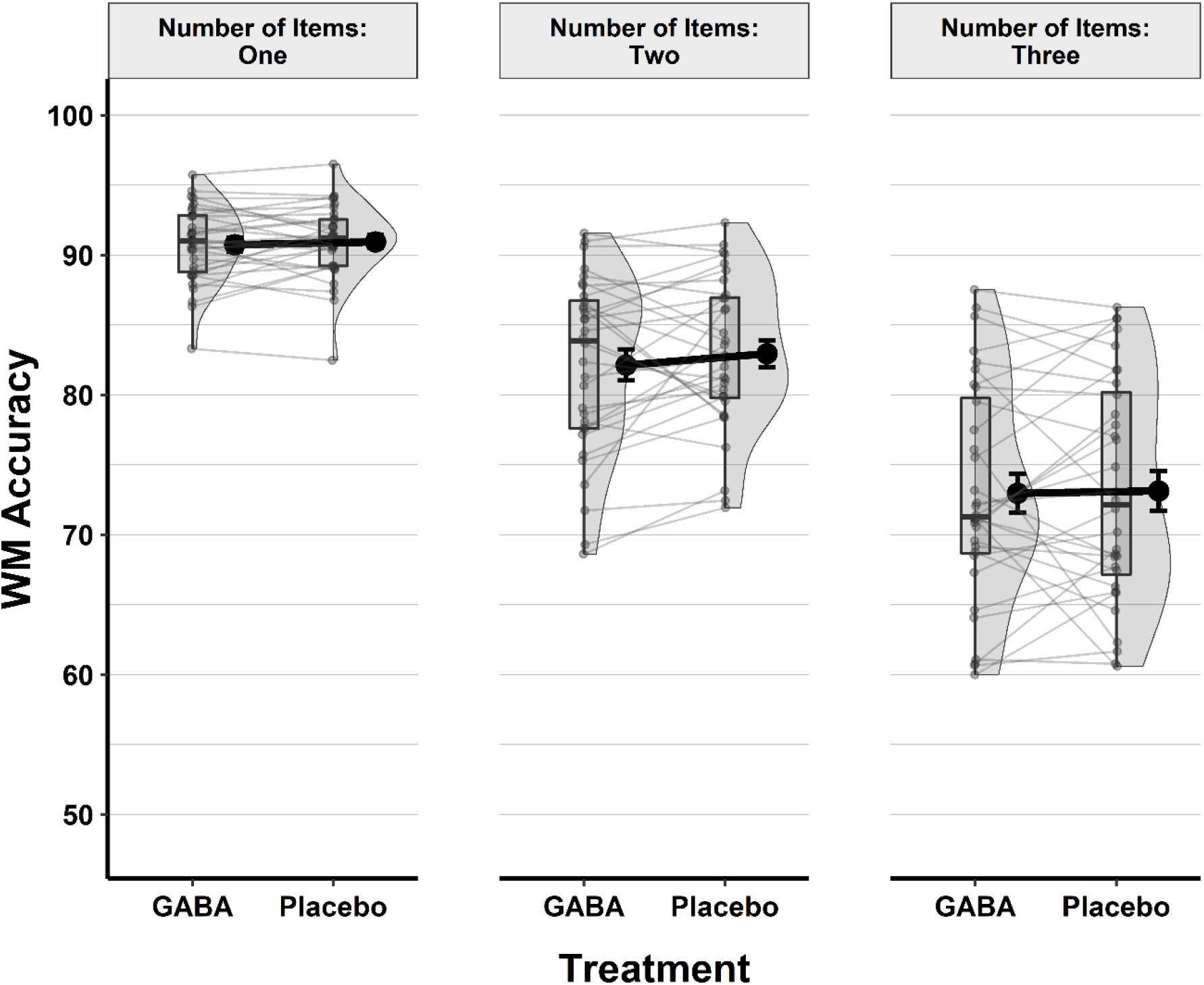
Accuracy in the Visual Working Memory task (percentage correct) by Treatment (GABA and Placebo) and the number of items. Boxplots show quartiles, black dots with error bars show means and standard errors, individual data points are presented with grey points and lines between individual data points show changes in treatment.

In the post-hoc pairwise comparisons, Tukey test results showed that working memory accuracy was higher when there was only one item (*Estimated marginal mean =* 90.8, SE = .46), than both when there were two (*Estimated marginal mean =* 82.5, SE = .97), Z = 12.62, *p* < .001, and three items (*Estimated marginal mean =* 73.0, SE = 1.32), Z = 17.32, *p* < .001. Also, accuracy was higher in the two item condition than when there were three items, Z = 16.20, *p* < .001. Regarding sessions, as might be expected, working memory accuracy was higher in the second session (*Estimated marginal mean =* 83.0, SE = .88), than in the first session (*Estimated marginal mean =* 81.3, SE = .88), Z = 6.38, *p* < .001. Lastly, on working memory accuracy, neither the gender of participants [*X*^2^(1) = 2.954, *p* = .086, *BF*_*10*_ = .055], nor the BMI scores [*X*^2^(1) = 2.840, *p* = .092, *BF*_*10*_ = .014] improved the model at all.s

## DISCUSSION

In this crossover designed, randomised, placebo-controlled, double-blind, counterbalanced and gender-balanced study, the acute effects of 800mg GABA supplementation on visual working memory precision, as well as temporal and spatial attention in healthy young adults were tested. We found that 800mg GABA intake acutely increased visual search reaction time, but did not modulate visual search accuracy, working memory maintenance, or temporal attention in the tested group. The results of the current study contrast with our original expectations of the effects of GABA on working memory, and spatial and temporal attention, which were based on previous studies, but in part these outcomes are also consistent with recent research findings.

### Spatial attention

Regarding spatial attention, in the current study, we found that 800mg GABA ingestion negatively affected visual search time. In other words, GABA intake acutely increased response times for visual search. However, these effects were limited to response time, as GABA did not acutely influence visual search accuracy. Although research on the acute effects of GABA on cognitive functions is limited, the current results are coherent with previous findings. Primarily, these come from a study by Lim and Aquili (2021), who carried out a study of 48 healthy young adults (32 females and 16 males) and tested the acute effects of 800 mg GABA administration on cognitive flexibility, measured by both task switching and Stroop tasks. GABA intake negatively influenced cognitive flexibility - namely task-switching performance, although there were no acute effects on response inhibition, as tested with the Stroop task. The authors reasoned that the relationship between GABA intake and cognitive performance might be nonlinear, causing the detrimental effect. Since we used the same dosage in the current study, this might also apply here.

Furthermore, it is known that alcohol is an agonist for GABA_A_ receptors and there is a consistent conclusion about the role of GABAergic actions in alcoholism. Abroms and Fillmore (2004) reported that alcohol intake acutely increased visual reaction time in healthy young social drinkers. Hironaka (2020) also concluded that alcohol ingestion acutely affects Stroop task error rates and response times in humans. Studies carried out with different agonists for GABA in animals have also shown similar results. Wardak et al. (2004) tested the effects of suppressing neural activity in the lateral intraparietal cortex in monkeys by administering muscimol, which is an agonist for GABA_A_ receptors, during a visual search task. The intraparietal cortex, in monkeys as well as in humans, is associated with perceptual-motor coordination (Tunik et al., 2007), and visual search behaviour (Bisley et al., 2011; Eckstein, 2011). Wardak et al. (2004) reported that inactivation in the intraparietal cortex affects visual search in a negative way and increases response times. Similarly, Hembrook et al. (2012) showed that inactivation by muscimol in prefrontal cortex caused increasing visual search time in rats. These findings suggest that GABA ingestion might similarly affect visual search response time via influencing GABAergic activation in prefrontal and parietal cortexes.

Apart from the negative effects of GABA on visual search time, the current study revealed that 800 mg GABA intake did not affect visual search accuracy. Using the same dose of GABA, Leonte et al. (2018) reported that there were no acute effects on spatial attention, both with regard to accuracy and reaction time. Task differences may explain this discrepancy. Leonte et al. (2018) measured visual search by a task asking participants to detect a target item amongst non-target items. Visual search accuracy ranged from 93.8% to 96%, and visual search times were between 348 ms and 367 ms, depending on the task condition. However, in the current study, participants were required to find the target item and identify its orientation (0°, 90°, 180°, 270°). Visual search accuracy ranged between 74.2 % and 85.7 % while reaction times varied between 985 ms to 1091 ms. In summary, it can be stated that the acute effects of GABA on visual search might be task- and difficulty-dependent, however, further studies are likely needed to make consistent and conclusive statements about these effects.

### Temporal attention

On the acute effects of GABA on temporal attention, in contrast to the previous study by Leonte et al. (2018), the current study found that 800mg of GABA intake does not affect temporal attention in healthy young adults. Leonte et al. (2018) found that GABA had a small positive effect on temporal attention. In their study, Leonte et al. found that at Lag 3, T2|T1 accuracy was 5.7 % higher in the GABA condition compared to the placebo. In the current study, no effect was found at all, although numerically at least there was a trend in the expected direction, as in the GABA condition T2|T1 accuracy was about 2 % higher compared to the placebo. Task differences might again explain these different outcomes. In the study by Leonte and colleagues, an RSVP task was also used with Lag 1, Lag 3 and Lag 8, but without a time restriction. It was furthermore an integration-specific version of the task, where participants were asked to distinguish whether shapes that could be integrated with each other were shown first or second (Akyürek et al., 2012). In the current study, however, targets were numbers, and there was a time restriction for responses: Participants were asked to give their responses as soon as T2 appeared. Nevertheless, with regard to the attentional blink, in Leonte’s study, the difference between T2|T1 accuracy at Lag 8 and Lag 3 was 13.9 %, while in the current study a difference of 21.6 % was found. Thus, if anything, this might indicate that the present speeded RSVP task causes greater variance and could be more sensitive. In spite of this, we failed to replicate the Lag 3 effect.

Reviewing these mixed outcomes to date, considering the acute positive effects of 800mg GABA on temporal attention with integration tasks, as well as the negative effects on cognitive flexibility and visual search time, and null effects on spatial and temporal attention with speeded RSVP, it might be speculated that GABA might have an effect on relatively simple attention- or inhibition-related tasks, but not more complex ones. Still, further studies using different tasks and/or different cognitive functions are needed to fully understand the effects of GABA intake on attention as well as other cognitive functions.

Alternatively, another account of the discrepant findings may be the biological gender of the participants. The study by Leonte et al. (2018) was not gender-balanced. Leonte et al. (2018) carried out their study with 32 young adults, but only four of them were male, and 28 were female. By contrast, the present study was gender-balanced, suggesting that the null findings could be related to the male subgroup. Gender balance should be taken into account as it has the potential to affect GABA’s effects on the brain. Previous work has linked differences in GABA levels in the brain with gender differences in mood disorders (Bjork et al., 2001; Sanacora et al., 1999) and alcoholism (Lancaster, 1994; Marlene Oscar-Berman, 2013). Furthermore, Cosgrove et al. (2007) showed that in addition to global cerebral blood flow, in the occipital cortex, cortical GABA levels are higher in females than males. Finally, Pandya et al. (2019) reported that in the superior temporal gyrus, which is associated with speech recognition (Nourski et al., 2021; Shekhar et al., 2019), GABA levels are higher in females than in males. Nevertheless, these studies generally also suggested that GABA actions are largely unaffected by gender differences. Indeed, in the present analyses, gender was not a reliable moderator of GABA effects in any of our tasks, thus casting doubt on the importance of this factor. More generally it might be noted that, as a result of not being gender-balanced, results from recent randomised control trials are still far from consistent in identifying whether gender affects the acute effects of GABA on cognitive functions. The current study is one of the first to test the acute effects of GABA on cognitive functions with gender-balanced data.

### Working memory

In addition to studying the effects of GABA on temporal and spatial attention, the current study also studied the effects on working memory maintenance and found that, again contrary to expectations, it was not improved by 800mg GABA. Although our study is one of the first to test the acute effects of GABA on working memory, indirect evidence from previous studies suggested that GABAergic modulation and activation may play a crucial role in the working memory process, particularly indirectly (Ellis & Nathan, 2001; Rammsayer et al., 2000; Rao et al., 2000). Several studies have also focused on GABA levels in different brain areas and their associations with working memory in both animals and humans (Auger & Floresco, 2015; Bañuelos et al., 2014; Gasbarri & Pompili, 2014; Heaney & Kinney, 2016; Marsman et al., 2017; Michels et al., 2012; Prut et al., 2010; Yoon et al., 2016). However, in general, previous studies observed the effects of GABA levels in the brain and not the effects of oral administration of GABA; the indirect relationships indicated by these previous studies may not be comparable with the results of studies that test the effects of direct ingestion.

It is also conceivable that the null effect of GABA ingestion in the current study is related to the specific task we used, in which items were to be maintained until recall, which does not involve active access or updating of working memory contents. In this context, a recent meta-analysis of 32 studies assessing the acute effects of alcohol administration by Spinola et al. (2022) might be relevant, since alcohol is an agonist to GABA receptors (Koob, 2004; Paul, 2006; Ticku, 1990); namely, it acts in the same way as GABA in the brain. It was reported that acute administration of alcohol impairs working memory performance with small to medium effects size, but that these effects depend on the type of working memory task. The effects are particularly strong and significant on working memory updating (as in the N-Back task) and manipulation (as in the self-ordered pointing task -SPOT-), but there were no significant effects on working memory maintenance (the Sternberg task). A similar pattern may apply to GABA ingestion, but this awaits direct empirical verification.

## Limitations

The current study also has several limitations. Its sample consisted solely of university students; it is generally assumed that young people (i.e., typical students) have high cognitive functioning; hence, it might be more difficult to observe a noticeable improvement in young people compared to older people (Alenius et al., 2019; Baird et al., 2019). Also, we did not measure physiological indicators that are associated with cognitive functions, such as heart rate and heart rate variability. Measuring both cognitive and physiological variables might produce a more comprehensive understanding of the effects of GABA on cognitive functions. Another limitation was that the current study tested participants’ cognitive functions 45 minutes after they consumed GABA, but the tests were only conducted once in each session, whether GABA or placebo. It would be better to conduct multiple tests, at 30, 45 and/or 70-minute intervals after the GABA intake, which would provide the opportunity to observe the acute effects of GABA on cognitive functions over time. Lastly, in the current study, participants consumed a single dose (800mg) GABA drink. To test its dose-dependent effects more efficiently, it would be better to serve different-sized doses, for instance, small (30mg), medium (200mg) and high (800mg).

## Conclusion

Despite the aforementioned limitations, the present evidence from our crossover designed, randomised, double-blind, counterbalanced, and gender-balanced study produced a clear outcome: The acute administration of 800mg of GABA increases visual search time significantly, but it does not modulate temporal attention or working memory maintenance in healthy young adults.

